# Engineering Orthogonal Carbon Dissimilation: A Gluconate Bypass Platform for Robust Stationary-Phase Biomanufacturing

**DOI:** 10.64898/2025.12.23.696260

**Authors:** Utsuki Yano, Payel Sarkar, Michael D. Lynch

**Author notes:** to whom all correspondence should be addressed.

## Abstract

Two-stage bioprocesses which decouple cell growth from product synthesis are an attractive approach to biomanufacturing. However high levels of production in stationary phase cultures often suffer from a progressive decline in metabolism. We demonstrate that in *E. coli* pyruvate accumulation, an inevitable consequence of high-flux metabolism, acts as a major inhibitor of stationary-phase glucose uptake. We present a novel central metabolism to optimize stationary phase production, a gluconate-bypass, which circumvents this challenge by rerouting carbon flux around glycolysis. This redesign achieves two critical outcomes: first, it decouples glucose uptake from pyruvate inhibition; second, glucose oxidation intrinsically co-generates the NADPH cofactor. Validated using NADPH-dependent L-alanine as a representative model, the GBP creates a self-regulating host that achieved a record titer of 197 g/L with a 1.6-fold extension of production longevity. This work establishes the GBP as a generalizable platform for robust stationary phase biosynthesis.

## Introduction

Microbial fermentation has advanced rapidly, driven by innovations in synthetic biology and metabolic engineering.[1–3] However, industrial competitiveness of microbial fermentation is often undermined by the adaptability of living cells, where complex regulatory networks dynamically adjust metabolism for survival rather than production consistency. [4–6] Two-stage fermentation mitigates this by separating growth from production,[7–12] but we have identified a critical “metabolic friction” in this approach: the decline of glucose uptake rates (GUR) during the stationary phase. Traditional engineering of glycolysis to resolve GUR bottlenecks is hindered by the pathway’s intricate and pleiotropic regulation.

In this study, we pinpoint pyruvate accumulation as a specific molecular bottleneck that impairs GUR in strains dependent on the native PTS and glycolytic flux. We introduce a fundamentally orthogonal solution: the gluconate-bypass (GBP) central metabolism. By replacing PTS transport with a permease and implementing a synthetic oxidative route that converts glucose to gluconate, we create a chassis resistant to pyruvate-induced inhibition. Furthermore, this architecture intrinsically balances precursor and cofactor pools by coupling glucose oxidation to NADPH generation, minimizing the need for external dynamic control elements. Using L-alanine as a high-flux test case, we demonstrate that the GBP platform redefines the performance limits of two-stage fermentation, offering a scalable blueprint for diverse biosynthetic pathways requiring sustained carbon and redox flux.

## Results

### Two-stage dynamic metabolic control enables robust alanine production by improving carbon and NADPH flux

In our previous work, we demonstrated robustness of two-stage dynamic metabolic control to improve the biosynthesis of several chemicals including L-alanine.[10,11] In this approach, the first stage supports biomass accumulation without metabolite production, and upon phosphate depletion, cells transition to stationary-phase production, activating the production pathway while downregulating competing metabolic pathways **(Fig. 1A)**. [10,13,14] This regulation is achieved using synthetic metabolic valves that dynamically reduce key central metabolic enzymes via controlled proteolysis, CRISPR-based gene silencing, or both.[10,15,16] Three valves—denoted “F,” “G,” and “U”—were previously implemented to enhance alanine production from glucose.[13,17] The “F” valve involves dynamic reductions in enoyl-ACP reductase (FabI) levels. Reduction in enoyl-ACP reductase levels reduces concentrations of acyl-ACPs (which are known inhibitors of the membrane bound transhydrogenase (PntAB)), thereby increasing NADPH synthesis. [18]The “G” valve involves dynamic reduction in citrate synthase (GltA) levels, leading to reduced α-ketoglutarate levels and relieving α-ketoglutarate–mediated inhibition of PTS-dependent glucose uptake.[18] Finally, the “U” valve involves the dynamic reduction in soluble transhydrogenase (UdhA) activity leading to elevated NADPH pools. Combined, these valves enhanced pyruvate and NADPH flux, resulting in a 6-fold increase in alanine production (∼3.5 g/L in micro-scale cultures) and ∼100 g/L in 6-L fermenters **(Fig. 1B)**. Despite these improvements, maximal production was sustained for only ∼40 hours, with a peak rate observed only in the first several hours of stationary phase. This reduction in production rate as the stationary phase progressed was apparent in the production of L-alanine from glucose and was not observed to the same degree or at all when a similar approach was used to produce xylose-derived products, such as xylitol (>100 hours). [10] To investigate the limitations in sustained alanine production and glucose uptake rate, we analyzed byproducts and found that pyruvate, the immediate precursor to alanine, accumulates to levels >5 g/L as productivity declines **(Fig. 1B)**. We hypothesize that pyruvate accumulation was responsible for slowing L-alanine production by impairing glucose uptake and glycolytic flux.

**Figure 1.**
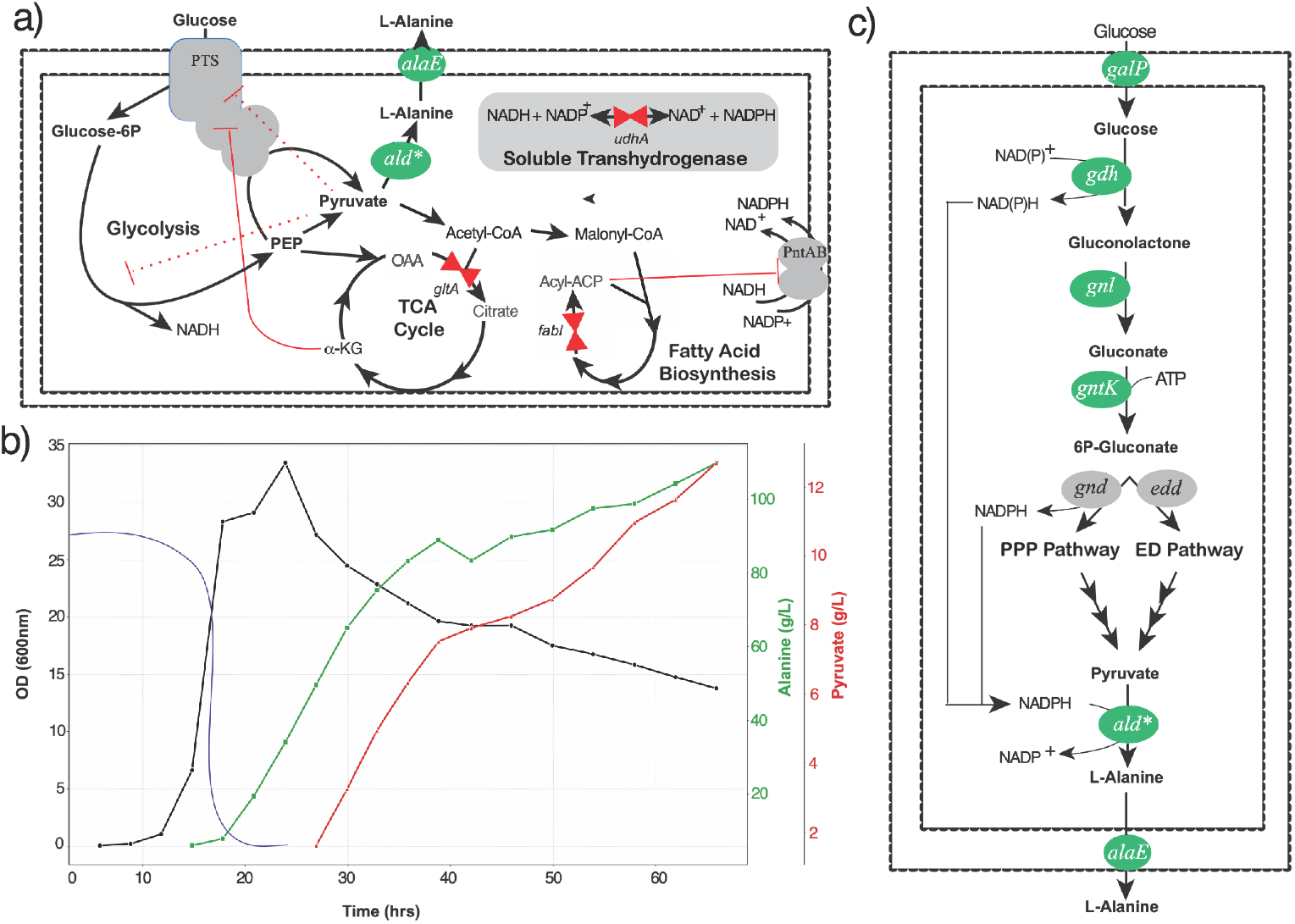
Overview of alanine production, previous production using PTS-dependent strain, and development of the gluconate-bypass system. (a) Overview of alanine production, highlighting the location of metabolic valves and key inhibitory metabolites within central metabolism. The alanine biosynthetic pathway is shown in green. Metabolic valves targeting citrate synthase (GltA), enoyl-ACP reductase (FabI), and soluble transhydrogenase (UdhA), which are dynamically controlled, are indicated by red triangles. (b) Alanine production in the PTS-dependent DLF_Z0025 strain containing the F, G, and U valves. Biomass is shown in black, alanine in green, pyruvate in red, and phosphate in blue. Phosphate levels are not measured but drawn to indicate the typical decline in phosphate concentration over time in these standard fermentations. (c) Design of the gluconate-bypass strain, in which glucose uptake via galactose permease (GalP) is oxidized to gluconolactone by glucose dehydrogenase (Gdh), hydrolyzed to gluconate by gluconolactonase (Gnl), and phosphorylated by gluconate kinase (GntK) to enter essential metabolic pathways.

### Pyruvate accumulation impairs glucose uptake rate in the stationary phase

To investigate this hypothesis, we conducted microfermentations with our “dynamic control” strain DLF_Z0025 **(Fig. 2A)**,[10] the starting strain used for L-alanine production lacking any metabolic valves. This strain relies on the native phosphotransferase system (PTS) for glucose uptake. Experiments were performed to spike pyruvate into the culture during the stationary phase. Without pyruvate, stationary phase GUR reached ∼0.45 g glucose/gCDW·h **(Fig. 2B)**. Introducing pyruvate at 10, 20, or 30 g/L reduced GUR by ∼50%, to ∼0.2 g glucose/gCDW·h. Next we spiked pyruvate into stationary phase cultures of this strain bearing plasmid pSMART-ala10, which enables L-alanine production.[11] Without pyruvate, alanine was produced at a specific rate of ∼0.12 g/gCDW·h, which decreased with pyruvate concentrations above 20 g/L **(Fig. 2C)**. These results support our hypothesis that pyruvate accumulation is responsible for reduction in GUR and L-alanine production rates in the PTS-dependent L-alanine production strain, although the exact mechanism has yet to be elucidated.

**Figure 2.**
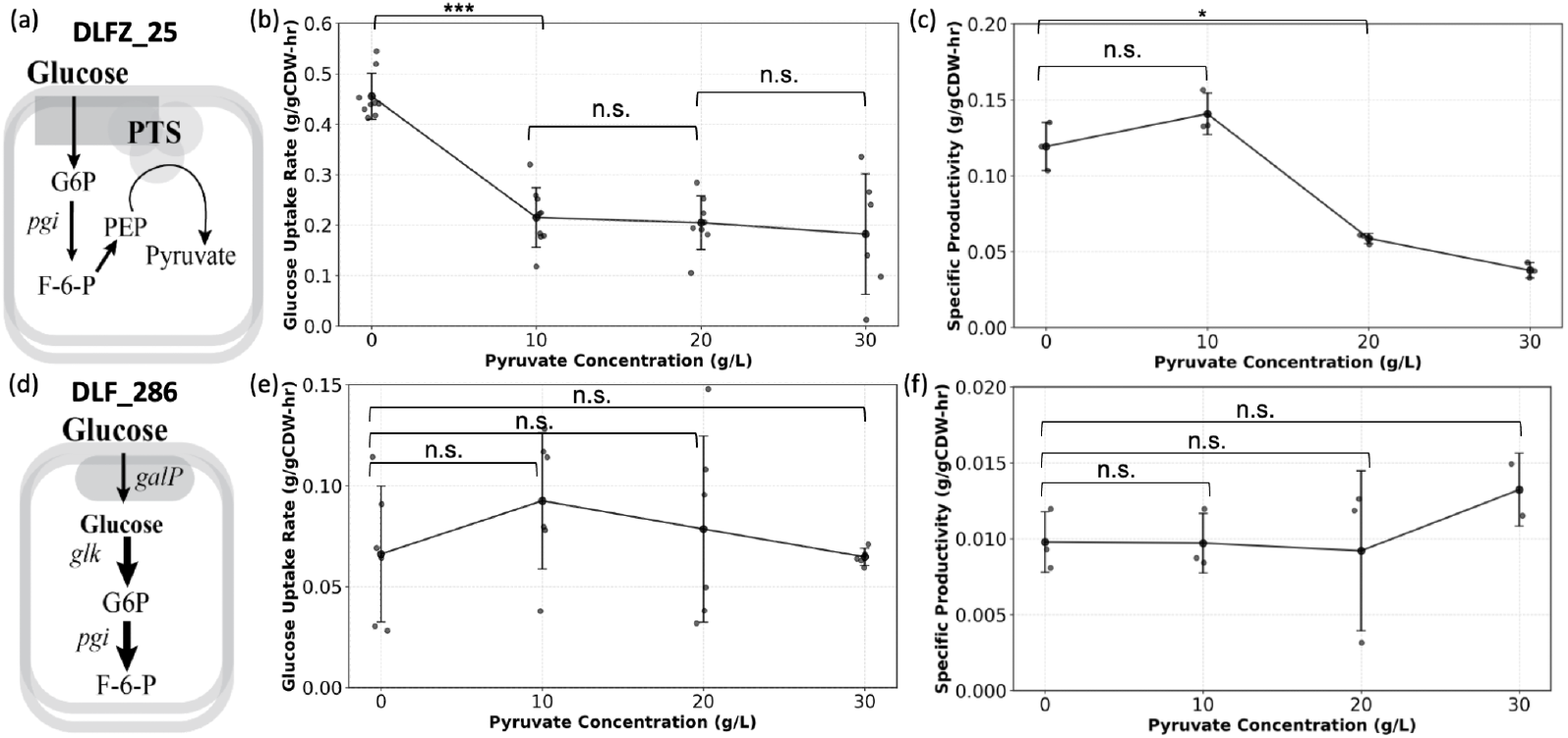
Phosphate-depleted stationary-phase glucose uptake and alanine productivity as a function of pyruvate concentration. (a) Design of strain DLF_Z0025. (b) Glucose uptake rate of DLF_Z0025 across different pyruvate concentrations. (c) Alanine productivity of DLF_Z0025 harboring the pSMART-ala10 plasmid. (d) Design of strain DLF_0286. (e) Glucose uptake rate of DLF_0286 across different pyruvate concentrations. (f) Alanine productivity of DLF_0286 harboring the pSMART-ala10 plasmid.

We next sought to see if the effect of pyruvate could be minimized by altering the glucose transport mechanism. To accomplish this we first constructed a PTS-independent strain (DLF_0286), as illustrated in **Figure 2D**. This was done by knocking out *ptsG*, which encodes the glucose-specific component of the PTS system, and replacing it with a codon-optimized, constitutively expressed glucokinase (glk), while also overexpressing the galactose permease (GalP). GalP has also been shown to be capable of glucose uptake.[19] We then repeated the stationary phase spike-in experiments that were performed for the PTS-dependent strain. Although the stationary phase GUR in this strain was unaffected by pyruvate accumulation, stationary phase GUR remained low even in the absence of pyruvate **(Fig. 2E)**. This low GUR limited alanine production, which remained minimal across different pyruvate concentrations **(Fig. 2F)**.

### Construction and characterization of strains with gluconate-bypass (GBP) central metabolism

In parallel with our construction of strain DLF_0286, we also sought to construct a strain that not only used an alternative sugar transport mechanism, but also bypasses glycolysis. Toward this goal we developed a strain with a gluconate-bypass central metabolism as illustrated in **Figure 1C**, which leverages not only an alternative glucose transport system, but additionally a glucose dehydrogenase which in combination with a lactonase can convert glucose to gluconate which can be metabolized via either pentose phosphate pathway or Entner-Duordoroff (ED) pathway in *E. coli*. Notably, similar oxidative glucose dissimilation pathways are naturally found in several microorganisms. In *Pseudomonas putida*, glucose is metabolized exclusively via the ED pathway through sequential periplasmic oxidations by glucose and gluconate dehydrogenases, yielding 2-ketogluconate that is subsequently converted to 6-phosphogluconate, the key intermediate linking the ED and pentose phosphate pathways.[20]^,[21]^ *Bacillus subtilis* utilizes cytosolic NAD(P)^+^-dependent glucose dehydrogenase and gluconolactonase to form gluconate, which feeds into the pentose phosphate pathway as an auxiliary oxidative route alongside glycolysis.[22] Importantly this design not only bypasses the natural *E. coli* metabolism we hypothesized to be inhibited by pyruvate, but also by using a NAD(P)^+^-dependent glucose dehydrogenase would allow for greater NADPH production.[23,24] In effect this new metabolic design was hypothesized to improve alanine production in theory by i) overcoming pyruvate inhibition, ii) alleviating the need to reduce citrate synthase (GltA) levels (which α-ketoglutarate pools that inhibit PTS-dependent glucose uptake) by employing a PTS-independent transport system, and iii) minimizing reliance on the *fabI* and *udhA* valves through increased NADPH formation via glucose oxidation.

This “Gluconate Bypass” (GBP) strain was constructed from the previously developed DLFS_0025 strain, which supports two-stage dynamic control.[25] This strain is isogenic to DLFZ_0025, but has modifications to increase the stability of silencing guide RNAs. The Gluconate bypass metabolism was implemented in DLFS_0025 by knocking out the native gntR, a repressor of gluconate kinase expression (*gntK)* and gluconate metabolism, and replacing it with Gdh, encoding a NAD(P)^+^-dependent glucose dehydrogenase, from *Bacillus cereus* and gluconolactonase (Gnl) from *Zymomonas mobilis* (**Fig. 3A)**.[26,27]^,[28]^ Together, these enzymes enable the oxidation of glucose to gluconate while generating either NADPH or NADH. Additionally, glucose transport was modified by deleting *ptsI/crr* and introducing the galactose permease (*galP*) (**Fig. 3B)**. After strain construction via recombineering,[29] multiple clones were screened for colony-to-colony variation (**Supplemental Figure S2**). Colony growth was compared to DLFZ_0025 and DLF_0286 in minimal media.[30] All clones exhibited consistent performance, showing ∼1.1-fold and ∼2.3-fold higher biomass after 24 h relative to DLFZ_0025 and DLF_0286, respectively (**Fig. 3D**). The expression from low phosphate inducible promoters [14] was then confirmed by measuring GFP expression upon phosphate depletion in microfermentations,[13] which was consistent across colonies and comparable to the prior two-stage host DLFZ_0025 (**Fig. 3D**). Based on these results, one isolate was selected for further characterization. Subsequently this strain was evaluated in a 1-L bioreactor, where growth in minimal media, glucose uptake, and phosphate consumption were compared to prior strains (**Fig. 3E**). Compared to the parental DLF_R002 strain, from which DLFZ_0025 was derived and is otherwise isogenic except for the deletion of the native sspB gene (enabling controlled proteolysis of metabolic valves), the GBP strain exhibited a maximum growth rate of 0.42 h^−1^ and a biomass yield of 0.29 g/g glucose, both comparable to those of DLF_R002 parent, which had a maximum growth rate of 0.41 h^−1^ and and a biomass yield of 0.34 g/g glucose.[13]

**Figure 3.**
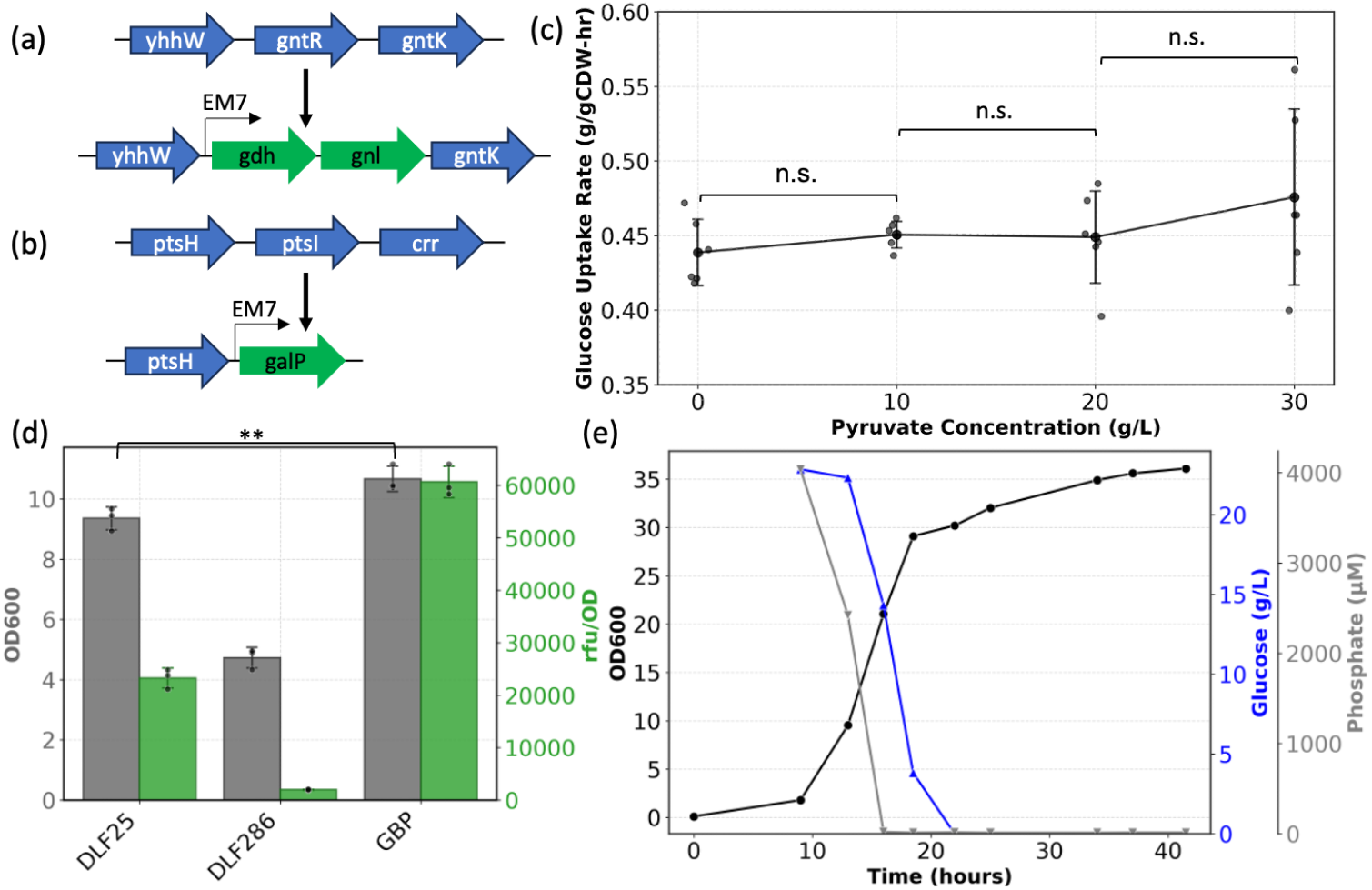
An overview of strain DLF_GBP1, engineered with a gluconate bypass of glycolysis. (a and b) Chromosomal modifications introduced to construct the gluconate-bypass (DLF_GBP1) strain. (a) Deletion of *gntR* and replacement with constitutively expressed *gdh* and *gnl*. (b) Deletion of *ptsI/crr* and replacement with constitutively expressed *galP*. (c) Glucose uptake rate of the DLF_GBP1 strain across different pyruvate concentrations. (d) Growth profiles and GFP expression of DLFZ_25, DLF_286, and DLF_GBP1 in 96-well microplates (e)Growth profile of the DLF_GBP1 strain and corresponding phosphate and glucose consumption in a 1-L bioreactor.

### GBP metabolism enable sustained GUR and alanine production even with pyruvate accumulation

Next we repeated the small scale spike-in studies to assess GUR and alanine production in the strains with the GBP metabolism in the presence of pyruvate. These strains maintained a GUR of 0.45 g/gCDW/h without pyruvate spike-in, comparable to DLFZ_0025. Upon introducing pyruvate (10, 20, 30 g/L), GUR remained unaffected **(Fig. 3C**). In fact, GUR consistently stayed above 0.45 g/gCDW/h, confirming that the new strain successfully overcomes the pyruvate-induced inhibition of glucose uptake and metabolism observed in strains reliant on glycolysis and PTS dependent transport. This sustained GUR translated to maintained alanine production rates in the presence of pyruvate (**Fig. 4A**). Notably, alanine production even increased with the addition of extracellular pyruvate, which we hypothesize is due to pyruvate uptake and conversion to L-alanine.[31,32]

**Figure 4.**
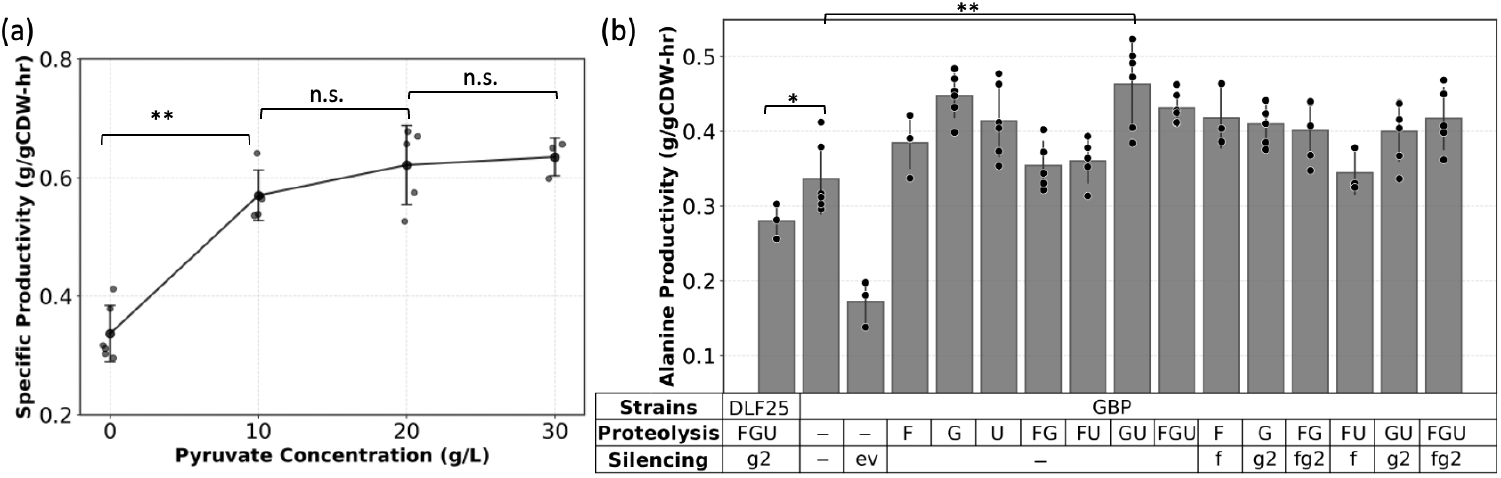
Alanine production using the gluconate-bypass (GBP) strain. (a) Alanine productivity of the strain across different pyruvate concentrations (b) Results of two-stage microfermentations producing alanine using different combination of “F”, “G”, “U” metabolic valves

### GBP strain possess optimal NADPH and pyruvate flux for efficient alanine production

Prior work with strain DLFZ_0025 demonstrated that introducing the “F,” “G,” and “U” valves was essential for achieving high alanine productivity by improving stationary phase sugar transport and enhancing NADPH supply. To evaluate whether these control elements remained beneficial in the gluconate-bypass (GBP) background, which features a restructured central metabolism, the same valve combinations were tested. Surprisingly, microfermentation results showed that the “G,” “U,” “GU,” and “FGU” variants all exhibited higher productivity than the no-valve control but were statistically indistinguishable from each other **(Fig. 4B)**. In contrast to DLFZ_0025, where all three valves were required, these GBP variants achieved comparable alanine productivities of approximately 0.4 g gCDW^−1^ h^−1^, about a 2-fold improvement over the DLFZ_0025 “FGU” benchmark. This suggests that the “F” valve, which boosts NADPH flux, is less critical in strains with GBP, as the initial oxidation of glucose to gluconolactone provides additional NADPH. Low cell density scale-up experiments in a 1-L bioreactors confirmed these findings: the DLF_GBP7 strain (“GU” valve strain) reached ∼123 g L^−1^ alanine after ∼65 h of production, with an average productivity of 0.28 g gCDW^−1^ h^−1^ **(Fig. 5A)**. The overall yield was 0.82 g alanine/g glucose, and the yield during the production phase reached 0.94 g alanine/g glucose. Importantly, production time was extended by ∼1.6-fold, and yield and titer both improved 1.2-fold with 43% less biomass, indicating more efficient carbon partitioning toward product formation. The GBP-GU strain also accumulated ∼60% less pyruvate than the previous DLFZ_0025 FGU strain, despite employing fewer dynamic control elements to enhance NADPH flux.

**Figure 5.**
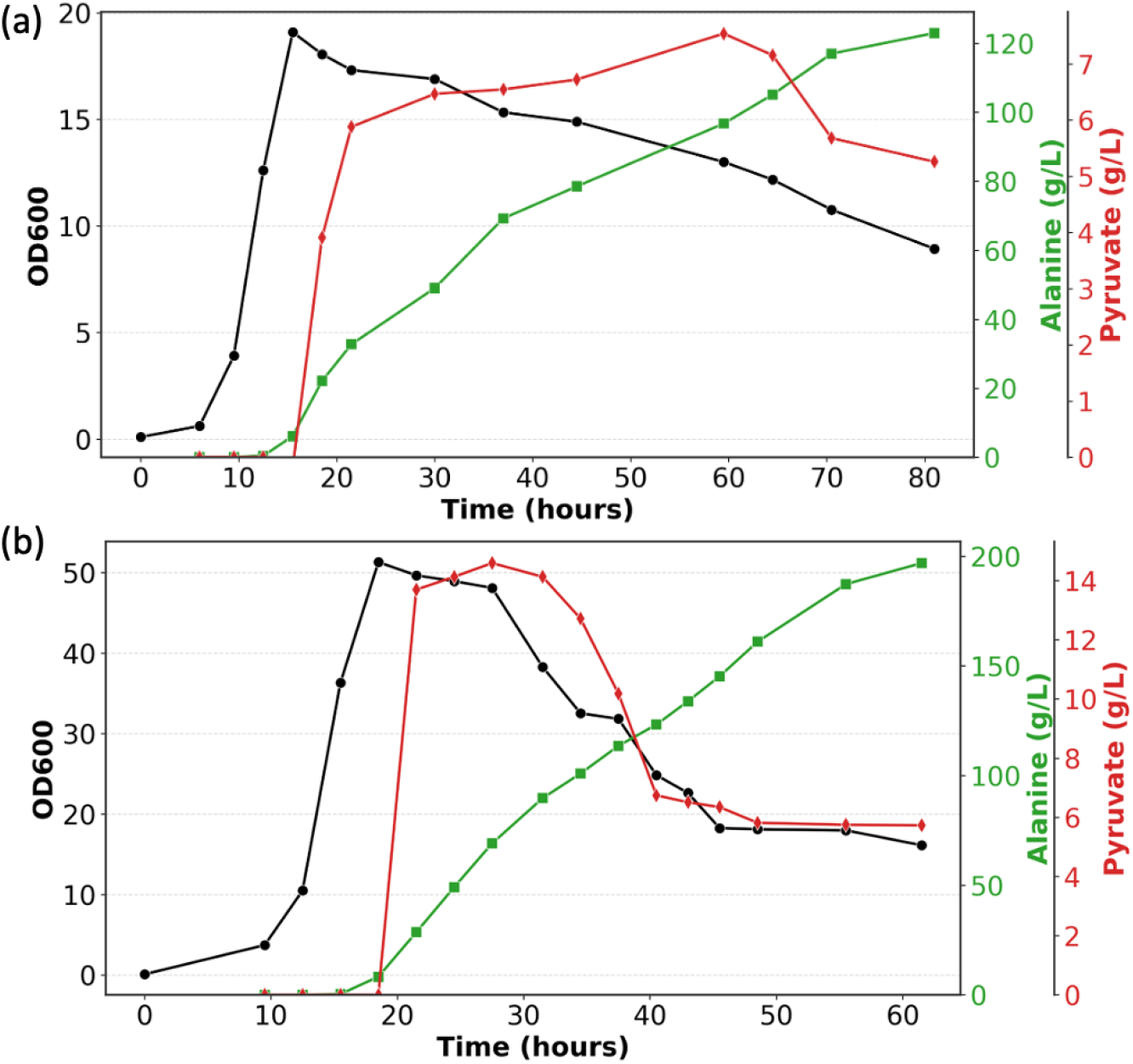
L-alanine production in 1-L bioreactors using the GU valve strain (DLF_GBP7) bearing pSMART-ala10. (a) Fermentation performed in FGM10 minimal media. Biomass (black lines), alanine titers (green lines), and pyruvate accumulation (red lines) were measured over time. (b) Fermentations performed in FGM25 minimal media. Biomass (black), alanine (green), and pyruvate (red) are shown.

Next we evaluated an intensified bioprocess with higher yet reasonable cell densities (∼18 gCDW/L) . Under these conditions strains DLF_GBP7 produced ∼197 g L^−1^ of alanine after ∼46 h of production, which is the highest reported titer to date demonstrated in *E. coli* **(Fig. 5B)**. This process had an average productivity of 0.24 g gCDW^−1^ h^−1^, overall yield was 0.81 g alanine/g glucose, and the yield during the production phase reached 0.92 g alanine/g glucose. Collectively, these results indicate that the gluconate-bypass central metabolism not only enables prolonged stationary phase production but also intrinsically helps to balance redox and carbon fluxes to sustain high productivity under production conditions.

## Discussion

Two-stage fermentations including those optimized with dynamic control offer significant promise in the pursuit of robust, high-titer bioprocesses. However, maintaining optimal metabolism and product synthesis in non-growing stationary phase cultures represents a critical challenge. Previous approaches to resolve this issue by directly manipulating glycolysis have been met with only limited success, reflecting the pathway’s intricate, essential, and highly regulated nature.[33–36] Our work first pinpoints and validates a critical, previously unrecognized bottleneck: pyruvate accumulation representing a major barrier to sustaining stationary-phase glucose uptake in *E. coli*. Pyruvate accumulations significantly impairs GUR, leading to reduced carbon flux and declining productivity. To overcome this imitation, we introduced a fundamentally different central metabolism to *E. coli*: the gluconate-bypass. This design couples PTS-independent transport with a synthetic pathway converting glucose to gluconate, thereby rerouting carbon flux entirely around the tightly regulated glycolytic nodes. The GBP architecture’s success is rooted in two synergistic effects. The gluconate-bypass enables high GUR and alanine production throughout the stationary phase, even under high concentrations of supplemented extracellular pyruvate (10, 20, and 30 g/L) Furthermore, the GBP architecture intrinsically integrates an essential co-factor generation step. Glucose oxidation inherently supplies the necessary NADPH for the conversion of pyruvate to L-alanine, eliminating the need for additional redox-balancing strategies. Together, these features enabled improved L-alanine production relative to traditional glycolysis-dependent strains, with alanine serving as a representative test case. GBP strains achieved the highest L-alanine titer reported to date in engineered *E. coli* grown on glucose with a significantly extended production phase. This demonstrated the utility of this novel central metabolic architecture for two-stage biosynthetic pathways that require sustained carbon flux and high NAD(P)H availability.

While strain DLF_GBP7 produced high levels of alanine and less pyruvate than prior strains, further improvements to NADPH regeneration could further enhance production by fully assimilating unconverted pyruvate early in the stationary phase to further improve production rates. Additionally, the modest biomass levels could also be increased further to enhance rates and titers. As titers continue to rise, optimizing analytical methods near the solubility limits of L-alanine are also critical. L-alanine is only soluble to ∼166 g L^−1^ at 25 °C and predicted to be about 178 g L^−1^ at 37 °C so adequate mixing and dilution is required for accurate quantification.[37,38] Moreover, L-alanine production is known to be more efficient under anaerobic conditions,[39] suggesting that adapting the GBP strain and optimizing process conditions for anaerobic fermentation could further improve productivity and yield.

In this work, we used L-alanine as a representative test case to demonstrate the architectural advantages of the gluconate-bypass (GBP) platform. By deregulating central metabolism from native glycolytic control, GBP sustains carbon flux during the production phase without inhibitory feedback while simultaneously enhancing NADPH availability through glucose oxidation. These coupled features enable robust two-stage production and are broadly advantageous for biosynthetic pathways that require sustained carbon flux and high reducing power, making this architecture extensible to a wide range of chemical products beyond L-alanine.

Traditional bioproduction strategies in engineered *E. coli* are intrinsically constrained by native regulatory networks, which impose fundamental limits on carbon flux and metabolic performance and thereby define the practical boundaries within which most strain engineering efforts operate. While two-stage production processes partially relax these constraints by decoupling growth from production, they remain subject to regulatory bottlenecks embedded within native central metabolism.

This work highlights the potential of constructing fundamentally different central metabolic architectures that are intrinsically less constrained by native regulation and are instead optimized for multi-stage processes and stationary-phase production. By introducing a gluconate-bypass central metabolism that is deregulated from canonical glycolytic control, we demonstrate that rerouting flux around tightly regulated nodes can yield robust, scalable, and self-regulating production phenotypes. L-alanine served as a representative test case and achieved the highest titer reported to date in engineered *E. coli* grown on glucose. This result demonstrates the general utility of the approach beyond a single product. More broadly, introducing deregulated central metabolism redefines regulatory boundaries and enables new design space for two-stage bioproduction across diverse chemical products.

## Methods

### Reagents and media

Unless otherwise specified, all materials and reagents were purchased from Sigma-Aldrich (St. Louis, MO, USA). Luria Broth (Lennox formulation) was used for routine strain and plasmid propagation and construction. Working antibiotic concentrations were as follows: kanamycin (35 μg/mL), chloramphenicol (35 μg/mL), zeocin (50 μg/mL), gentamicin (25 μg/mL), blasticidin (100 μg/mL), and spectinomycin (50 μg/mL). SM10++, SM10 (no phosphate), FGM10, and FGM25 media were prepared according to previously reported protocols[13,14].

### Plasmids and strains

All host strains were constructed using standard recombineering methods as previously described.[29] A complete list of strains is provided in Supplementary Table S2. The recombineering plasmid pSIM5 was kindly provided by Donald Court (NCI; https://redrecombineering.ncifcrf.gov/court-lab.html).[40] Recombinant strains were verified by PCR and DNA sequencing (Azenta Life Sciences, MA).

Alanine production plasmids were constructed using synthetic DNA fragments cloned into the pSMART-HCKan vector (Lucigen, WI). Plasmid assembly was carried out using the NEBuilder® HiFi DNA Assembly Master Mix according to the manufacturer’s instructions (New England Biolabs, MA). All plasmid sequences were confirmed by DNA sequencing (Azenta Life Sciences, MA) and deposited with Addgene. Primers and gBlocks used in this study are listed in Supplementary Table S3 and S4.

### Seed Preparation

A 5 mL Luria Broth (LB) culture supplemented with the appropriate antibiotics was inoculated from glycerol stock and incubated overnight at 37 °C with shaking. The following day, 1% (v/v) of the overnight culture was transferred into a 250 mL baffled shake flask containing 50 mL SM10++ media with antibiotics and incubated at 37 °C until the culture reached an OD_600_ of 6–10. Subsequently, 1% (v/v) of this culture was used to inoculate 50 mL of SM10 media (containing half the concentrations of casamino acids and yeast extract present in SM10++), and incubation was continued under identical conditions until OD_600_ reached 6–10. Finally, 1% (v/v) of this culture was used to inoculate SM10 media lacking casamino acids and yeast extract, and cells were harvested when OD_600_ again reached 6–10.

Cells were collected by centrifugation at 4,000 rpm for 15 min, and the resulting pellet was resuspended in FGM10 media to an OD_600_ of 10. Seed vials were prepared by mixing 6.5 mL of the resuspended culture with 1.5 mL sterile 50% (v/v) glycerol and stored at −70 °C.

### Fermentations

Minimal media microfermentations were performed as previously described.[14] For pyruvate spike-in experiments, sodium pyruvate was added after the wash step to final concentrations of 10, 20, or 30 g L^−1^. Instrumented bioreactor fermentations were conducted following the protocol of Menacho-Melgar et al. (2020), with minor modifications to the glucose feeding strategy to ensure adequate carbon supply. For low cell density fermentation using FGM10 media, tanks were filled with 800 mL of FGM10 media formulation as previously reported containing an appropriate phosphate concentration to achieve a final *E. coli* biomass of approximately 10 gCDW/L and the starting glucose concentration of 35 g L^−1^.[13] Antibiotics were added as needed. Frozen seed vials containing 8 mL of seed culture were used to inoculate the tanks. Temperature and pH within the bioreactors were controlled at 37 °C and 6.8, respectively, using 10 M ammonium hydroxide and 1 M hydrochloric acid as titrants. The glucose feed concentration was 500 g L^−1^, and the feed rate was maintained at 7 g h^−1^ for the first 18.5 h, reduced to 0.75 g h^−1^ for the subsequent 11.5 h, paused for 14.5 h, and then resumed at 0.75 g h^−1^ for an additional 21.5 h.

For medium cell density fermentation using FGM25 media, tanks were filled with 600 mL of FGM25 media formulation as previously reported containing an appropriate phosphate concentration to achieve a final *E. coli* biomass of approximately 25 gCDW L^−1^.[13] Antibiotics were added as needed. The initial media glucose concentration for this run was 55 g L^−1^. Concentrated sterile filtered glucose feed (700 g L^−1^) was added to the tanks at an initial rate of 8.25 g h^−1^ when the rate of agitation increased above 1000 rpm. This rate was then increased exponentially from 8.25 g h^−1^ to 35.7 g h^−1^, until 60 g total glucose had been added, at this point the feed was set to 21 g h^−1^ for 3 hours and then maintained at 15.6 g h^−1^ for 6 hours to maximize L-alanine production and finally reduced to 10.4 g h^−1^ until the remaining glucose was fed inside the tank.

Average productivity was calculated by dividing the total alanine produced (g) by the production time and maximum biomass concentration. Production yield was determined by first calculating the total glucose consumed during the production phase, subtracting the amount of glucose converted into alanine prior to that phase, and then dividing the alanine produced by the corrected glucose consumption.

### Expression Analysis & Densitometry

Expression of pSMART-ala10 and pSMART-ala10NADH was carried out in the DLFZ_25 strain using the microtiter plate fermentation protocol. Protein expression levels were quantified by densitometric analysis of SDS–PAGE gels. Whole-cell samples were normalized to OD600=10, mixed 1:1 with 2× loading buffer, and heated at 95 °C for 10 min. A 10 µL aliquot of each sample was loaded onto a 4–15% gradient Mini-PROTEAN TGX precast gel (Bio-Rad Laboratories, CA) and ran at 160 V. Densitometry was performed using FIJI.[41,42]

### Analytical methods

L-Alanine concentrations were determined as previously described.[11]

Glucose concentrations in microfermentation and bioreactor samples were measured using a UPLC system (Acquity H-Class, Waters Corp., MA, USA) equipped with a 2414 Refractive Index (RI) detector. Chromatographic separation was achieved on a Rezex ROA-Organic Acid H^**+**^ (8%) column (300 × 7.8 mm; Cat. no. 00H-0138-K0, Phenomenex, Inc., CA, USA) maintained at 50 °C. The mobile phase consisted of 5 mM sulfuric acid, delivered isocratically at 0.5 mL min^−1^ for 35 min. The injection volume was 10 μL. To maintain linearity within the analytical range, samples were diluted 5-fold (microfermentation) or 10–20-fold (bioreactor) with deionized water prior to analysis. For glucose quantification in pyruvate spike-in samples, an alternative UPLC-RI method was employed to avoid co-elution of glucose and pyruvate. Separation was performed using an Acquity BEH Amide column (2.1 × 150 mm, 1.7 μm; Cat. no. 186004802, Waters Corp., MA, USA) maintained at 65 °C. The mobile phase consisted of 80% (v/v) acetonitrile and 20% (v/v) water containing 5 mM ammonium formate and 0.06% (v/v) ammonium hydroxide, delivered isocratically at 0.75 mL min^−1^ for 15 min.

Pyruvate concentrations were determined using the same HPLC method described above with the Rezex ROA-Organic Acid H^**+**^ (8%) column, coupled to a Waters TUV detector (Waters Corp., MA, USA). Absorbance was monitored at 210 nm. Samples were diluted 20-fold (microfermentation) or 200-fold (bioreactor) with deionized water prior to analysis to maintain linearity.

Phosphate concentration was determined using a Phosphate Assay Kit (Sigma-Aldrich, St. Louis, MO, USA) following the manufacturer’s instructions. Briefly, culture samples were centrifuged to remove cells, and the resulting supernatant was used for the assay.

## Author contributions

U. Yano and P. Sarkar constructed plasmids and strains, performed microfermentations and instrumented fermentations and analytical analyses. M.D. Lynch designed experiments. All authors analyzed results, wrote, revised and edited the manuscript.

## Conflicts of Interest

M.D. Lynch has a financial interest in DMC Biotechnologies, Inc., Roke Biotechnologies, LLC, and DINYA DNA, Inc

## Data Availability

All Data is made available in the manuscript or supplemental Materials

## Acknowledgements

We would like to acknowledge the following support: DOE EE0007563, NSF # 2350533, U. Yano was in part supported by the Takenaka Scholarship Foundation.

## References

1. Cameron, D.E. et al. (2014) A brief history of synthetic biology. Nat. Rev. Microbiol. 12, 381–390

2. Carruthers, D.N. and Lee, T.S. (2022) Translating advances in microbial bioproduction to sustainable biotechnology. Front. Bioeng. Biotechnol. 10, 968437

3. Nielsen, J. and Keasling, J.D. (2016) Engineering cellular metabolism. Cell 164, 1185–1197

4. Wehrs, M. et al. (2019) Engineering robust production microbes for large-scale cultivation. Trends Microbiol. 27, 524–537

5. Olsson, L. et al. (2022) Robustness: linking strain design to viable bioprocesses. Trends Biotechnol. 40, 918–931

6. Adegboye, M.F. et al. (2021) Bioprospecting of microbial strains for biofuel production: metabolic engineering, applications, and challenges. Biotechnol. Biofuels 14, 5

7. Burg, J.M. et al. (2016) Large-scale bioprocess competitiveness: the potential of dynamic metabolic control in two-stage fermentations. Curr. Opin. Chem. Eng. 14, 121–136

8. Chubukov, V. and Sauer, U. (2014) Environmental dependence of stationary-phase metabolism in Bacillus subtilis and Escherichia coli. Appl. Environ. Microbiol. 80, 2901–2909

9. Kolter, R. et al. (1993) The stationary phase of the bacterial life cycle. Annu. Rev. Microbiol. 47, 855–874

10. Li, S. et al. (2021) Dynamic control over feedback regulatory mechanisms improves NADPH flux and xylitol biosynthesis in engineered E. coli. Metab. Eng. 64, 26–40

11. Ye, Z. et al. (2021) Two-stage dynamic deregulation of metabolism improves process robustness & scalability in engineered E. coli. Metab. Eng. 68, 106–118

12. Hennigan, J.N. et al. (2024) Scalable, robust, high-throughput expression & purification of nanobodies enabled by 2-stage dynamic control. Metab. Eng. 85, 116–130

13. Menacho-Melgar, R. et al. (2020) Scalable, two-stage, autoinduction of recombinant protein expression in E. coli utilizing phosphate depletion. Biotechnol. Bioeng. 117, 2715–2727

14. Moreb, E.A. et al. (2020) Media Robustness and Scalability of Phosphate Regulated Promoters Useful for Two-Stage Autoinduction in E. coli. ACS Synth. Biol. 9, 1483–1486

15. Luo, M.L. et al. (2015) Repurposing endogenous type I CRISPR-Cas systems for programmable gene repression. Nucleic Acids Res. 43, 674–681

16. McGinness, K.E. et al. (2006) Engineering controllable protein degradation. Mol. Cell 22, 701–707

17. Menacho-Melgar, R. et al. (2020) Improved two-stage protein expression and purification via autoinduction of both autolysis and auto DNA/RNA hydrolysis conferred by phage lysozyme and DNA/RNA endonuclease. Biotechnol. Bioeng. 117, 2852–2860

18. Li, S. et al. (2020) Dynamic control over feedback regulation identifies pyruvate-ferredoxin oxidoreductase as a central metabolic enzyme in stationary phase E. coli bioRxiv, 2020.07.26.219949

19. Hernández-Montalvo, V. et al. (2003) Expression of galP and glk in a Escherichia coli PTS mutant restores glucose transport and increases glycolytic flux to fermentation products. Biotechnol. Bioeng. 83, 687–694

20. del Castillo, T. et al. (2007) Convergent peripheral pathways catalyze initial glucose catabolism in Pseudomonas putida: genomic and flux analysis. J. Bacteriol. 189, 5142–5152

21. Rozova, O.N. et al. (2021) Enzymes of an alternative pathway of glucose metabolism in obligate methanotrophs. Sci. Rep. 11, 8795

22. Zamboni, N. et al. (2004) The Bacillus subtilis yqjI gene encodes the NADP+-dependent 6-P-gluconate dehydrogenase in the pentose phosphate pathway. J. Bacteriol. 186, 4528–4534

23. Weckbecker, A. and Hummel, W. (2005) Glucose dehydrogenase for the regeneration of NADPH and NADH. In Microbial Enzymes and Biotransformations, pp. 225–238, Humana Press

24. Wu, Y. et al. (2022) Constitutive glucose dehydrogenase elevates intracellular NADPH levels and luciferase luminescence in Bacillus subtilis. Microb. Cell Fact. 21, 266

25. Ye, Z. et al. (2021) Escherichia coli Cas1/2 Endonuclease Complex Modifies Self-Targeting CRISPR/Cascade Spacers Reducing Silencing Guide Stability. ACS Synth. Biol. 10, 29–37

26. Bach, J.A. and Sadoff, H.L. (1962) Aerobic sporulating bacteria i. J. Bacteriol. 83, 699–707

27. Goldman, M. and Blumenthal, H.J. (1964) Pathways of glucose catabolism in Bacillus cereus1. J. Bacteriol. 87, 377–386

28. Kanagasundaram, V. and Scopes, R. (1992) Isolation and characterization of the gene encoding gluconolactonase from Zymomonas mobilis. Biochim. Biophys. Acta 1171, 198–200

29. Li, X.-T. et al. (2013) Positive and negative selection using the tetA-sacB cassette: recombineering and P1 transduction in Escherichia coli. Nucleic Acids Res. 41, e204

30. Li, S. et al. (2024) 2-stage microfermentations. Metab. Eng. Commun. 18, e00233

31. Kristoficova, I. et al. (2018) BtsT, a Novel and Specific Pyruvate/H+ Symporter in Escherichia coli. J. Bacteriol. 200

32. Gasperotti, A. et al. (2020) Function and Regulation of the Pyruvate Transporter CstA in Escherichia coli. Int. J. Mol. Sci. 21, 9068

33. Chubukov, V. et al. (2017) Engineering glucose metabolism of Escherichia coli under nitrogen starvation. NPJ Syst. Biol. Appl. 3, 16035

34. Michalowski, A. et al. (2017) Escherichia coli HGT: Engineered for high glucose throughput even under slowly growing or resting conditions. Metab. Eng. 40, 93–103

35. Traxler, M.F. et al. (2008) The global, ppGpp-mediated stringent response to amino acid starvation in Escherichia coli. Mol. Microbiol. 68, 1128–1148

36. Kyselova, L. et al. (2018) Type and capacity of glucose transport influences succinate yield in two-stage cultivations. Microb. Cell Fact. 17, 132

37. Dunn, M.S. et al. (1933) The solubility of the amino acids in water. J. Biol. Chem. 103, 579–595

38. Chua, Y.Z. et al. (2018) New experimental melting properties as access for predicting amino-acid solubility. RSC Adv. 8, 6365–6372

39. Zhang, X. et al. (2007) Production of L-alanine by metabolically engineered Escherichia coli. Appl. Microbiol. Biotechnol. 77, 355–366

40. Sharan, S.K. et al. (2009) Recombineering: a homologous recombination-based method of genetic engineering. Nat. Protoc. 4, 206–223

41. Schindelin, J. et al. (2012) Fiji: an open-source platform for biological-image analysis. Nat. Methods 9, 676–682

42. Rueden, C.T. et al. (2017) ImageJ2: ImageJ for the next generation of scientific image data. BMC Bioinformatics 18, 529

