## Supplemental Materials for "Engineering Orthogonal Carbon Dissimilation: A Gluconate Bypass Platform for Robust Stationary-Phase Biomanufacturing"

**Table S1:** Plasmids used in this study

| Plasmid | Insert | Ori | Res | Addgene | Source |
|---|---|---|---|---|---|
| pSMART-HC-Kan | None | colE1 | Kan | N/A | Lucigen |
| pHCKan-yibDp-GFPuv | GFPuv | colE1 | Kan | 127078 | (Menacho-Melgar et al. 2020) |
| pSMART-Ala10 | yibDp-ald*-alaE | colE1 | Kan | 87135 | (Menacho-Melgar et al. 2020) |
| pSIM5 | Recombineering genes | pSC101ts | Cm | N/A | Court Lab |
| pCASCADE-ev | empty gRNA control | p15a | Cm | 65821 | (Li et al. 2020) |
| pCASCADE-f | fabIp silencing guide | p15a | Cm | 66635 | (Ye et al. 2021) |
| pCASCADE-g2 | gltA2p silencing gRNA | p15a | Cm | 65817 | (Li et al. 2020) |

**Table S2:** Strains used in this study

| Strain | Genotype | Source |
|---|---|---|
| DLF_Z0025 | F-, λ-, Δ(araD-araB)567, lacZ4787(del)(::rrnB-3) , rph-1, Δ(rhaD-rhaB)568, hsdR514, ΔackA-pta, ΔpoxB, ΔpflB, ΔldhA, ΔadhE, ΔiclR, ΔarcA, ΔsspB, Δcas3::tm-ugpb-sspB-pro-casA. | (Li et al. 2020) |
| DLF_S0025 | F-, λ-, Δ(araD-araB)567, lacZ4787(del)(::rrnB-3) , rph-1, Δ(rhaD-rhaB)568, hsdR514, ΔackA-pta, ΔpoxB, ΔpflB, ΔldhA, ΔadhE, ΔiclR, ΔarcA, ΔsspB::frt, Δcas3:: ugpBp-sspB-yibDp-casA | (Ye et al. 2021) |
| DLF_Z0047 | DLF_Z0025, fabI-DAS+4-gentR, gltA-DAS+4::zeoR, udhA-DAS+4-bsdR | (Li et al. 2021) |
| DLF_0286 | DLF_Z0025, ΔptsG::proCp-glk, galPp::proCp | This study |
| DLF_GBP1 | DLF_S0025, ΔptsI/crr::EM7-galP, ΔgntR::EM7-gdh-gnl | This study |
| DLF_GBP2 | DLF_GBP1, fabI-DAS+4-gentR | This study |
| DLF_GBP3 | DLF_GBP1, gltA-DAS+4-zeoR | This study |
| DLF_GBP4 | DLF_GBP1, udhA-DAS+4-bsdR | This study |
| DLF_GBP5 | DLF_GBP1, fabI-DAS+4-gentR, gltA-DAS+4-zeoR | This study |
| DLF_GBP6 | DLF_GBP1, fabI-DAS+4-gentR, udhA-DAS+4-bsdR | This study |

| | | |
|---|---|---|
| DLF_GBP7 | DLF_GBP1, gltA-DAS+4-zeoR, udhA-DAS+4-bsdR | This study |
| DLF_GBP8 | DLF_GBP1, fabI-DAS+4-gentR, gltA-DAS+4-zeoR, udhA-DAS+4-bsdR | This study |

**Table S3:** Synthetic DNA utilized for strain construction.

| |
|---|
| **fabI-DAS4-gentR** |
| CTATTGAAGATGTGGGTAACTCTGCGGCATTCCTGTGCTCCGATCTCTCTGCCGGTATCTCCGGTGAAGTGGT<br>CCACGTTGACGGCGGTTTCAGCATTGCTGCAATGAACGAACTCGAACTGAAAGCGGCCAACGATGAAAACTAT<br>TCTGAAAACTATGCGGATGCGTCTTAATAGGAAGTTCCTATTCTCTAGAAAGTATAGGAACTTCCGAATCCATGT<br>GGGAGTTTATTCTTGACACAGATATTTATGATATAATAACTGAGTAAGCTTAACATAAGGAGGAAAAACATATGTTA<br>CGCAGCAGCAACGATGTTACGCAGCAGGGCAGTCGCCCTAAAACAAAGTTAGGTGGCTCAAGTATGGGCATCA<br>TTCGCACATGTAGGCTCGGCCCTGACCAAGTCAAATCCATGCGGGCTGCTCTTGATCTTTTCGGTCGTGAGTT<br>CGGAGACGTAGCCACCTACTCCCAACATCAGCCGGACTCCGATTACCTCGGGAACTTGCTCCGTAGTAAGACA<br>TTCATCGCGCTTGCTGCCTTCGACCAAGAAGCGGTTGTTGGCGCTCTCGCGGCTTACGTTCTGCCCAAGTTTG<br>AGCAGCCGCGTAGTGAGATCTATATCTATGATCTCGCAGTCTCCGGCGAGCACCGGAGGCAGGGCATTGCCAC<br>CGCGCTCATCAATCTCCTCAAGCATGAGGCCAACGCGCTTGGTGCTTATGTGATCTACGTGCAAGCAGATTACG<br>GTGACGATCCCGCAGTGGCTCTCTATACAAAGTTGGGCATACGGGAAGAAGTGATGCACTTTGATATCGACCCA<br>AGTACCGCCACCTAAGAAGTTCCTATTCTCTAGAAAGTATAGGAACTTCCGTTCTGTTGGTAAAGATGGGCGGC<br>GTTCTGCCGCCCGTTATCTCTGTTATACCTTTCTGATATTTGTTATCGCCGATCCGTCTTTCTCCCCTTCCCGCC<br>TTGCGTCAGG |
| **udhA-DAS+4-bsdR** |
| TCTGGGTATTCACTGCTTTGGCGAGCGCGCTGCCGAAATTATTCATATCGGTCAGGCGATTATGGAACAGAAAG<br>GTGGCGGCAACACTATTGAGTACTTCGTCAACACCACCTTTAACTACCCGACGATGGCGGAAGCCTATCGGGT<br>AGCTGCGTTAAACGGTTTAAACCGCCTGTTTGCGGCCAACGATGAAAACTATTCTGAAAACTATGCGGATGCGT<br>CTTAATAGTTGACAATTAATCATCGGCATAGTATATCGGCATAGTATAATACGACTCACTATAGGAGGGCCATCATG<br>AAGACCTTCAACATCTCTCAGCAGGATCTGGAGCTGGTGGAGGTCGCCACTGAGAAGATCACCATGCTCTATG<br>AGGACAACAAGCACCATGTCGGGGCGGCCATCAGGACCAAGACTGGGGAGATCATCTCTGCTGTCCACATTG<br>AGGCCTACATTGGCAGGGTCACTGTCTGTGCTGAAGCCATTGCCATTGGGTCTGCTGTGAGCAACGGGCAGA<br>AGGACTTTGACACCATTGTGGCTGTCAGGCACCCCTACTCTGATGAGGTGGACAGATCCATCAGGGTGGTCAG<br>CCCCTGTGGCATGTGCAGAGAGCTCATCTCTGACTATGCTCCTGACTGCTTTGTGCTCATTGAGATGAATGGCA<br>AGCTGGTCAAAACCACCATTGAGGAACTCATCCCCCTCAAGTACACCAGGAACTAAAGTAAAACTTTATCGAAA<br>TGGCCATCCATTCTTGCGCGGATGGCCTCTGCCAGCTGCTCATAGCGGCTGCGCAGCGGTGAGCCAGGACGA<br>TAAACCAGGCCAATAGTGCGGCGTGGTTCCGGCTTAATGCACGG |
| **gltA-DAS+4-zeoR** |
| GTATTCCGTCTTCCATGTTCACCGTCATTTTCGCAATGGCACGTACCGTTGGCTGGATCGCCCACTGGAGCGA<br>AATGCACAGTGACGGTATGAAGATTGCCCGTCCGCGTCAGCTGTATACAGGATATGAAAAACGCGACTTTAAAA<br>GCGATATCAAGCGTGCGGCCAACGATGAAAACTATTCTGAAAACTATGCGGATGCGTCTTAATAGTTGACAATTA<br>ATCATCGGCATAGTATATCGGCATAGTATAATACGACTCACTATAGGAGGGCCATCATGGCCAAGTTGACCAGTG<br>CCGTTCCGGTGCTCACCGCGCGCGACGTCGCCGGAGCGGTCGAGTTCTGGACCGACCGGCTCGGGTTCTC<br>CCGGGACTTCGTGGAGGACGACTTCGCCGGTGTGGTCCGGGACGACGTGACCCTGTTCATCAGCGCGGTCC<br>AGGACCAGGTGGTGCCGGACAACACCCTGGCCTGGGTGTGGGTGCGCGGCCTGGACGAGCTGTACGCCGA |

GTGGTCGGAGGTCGTGTCCACGAACTTCCGGGACGCCTCCGGGCCGGCCATGACCGAGATCGGCGAGCAG
CCGTGGGGGCGGGAGTTCGCCCTGCGCGACCCGGCCGGCAACTGCGTGCACTTTGTGGCAGAGGAGCAGG
ACTGAGGATAAGTAATGGTTGATTGCTAAGTTGTAAATATTTTAACCCGCCGTTCATATGGCGGGTTGATTTTTAT
ATGCCTAAACACAAAAAATTGTAAAAATAAAATCCATTAACAGACCTATATAGATATTTAAAAAGAATAGAACAGCT
CAAATTATCAGCAACCCAATACTTTCAATTAAAAACTTCATGGTAGTCGCATTTATAACCCTATGAAA

**ΔptsI/crr::EM7-galP-specR**

TGTGACTATCTCCGCAGAAGGCGAAGACGAGCAGAAAGCGGTTGAACATCTGGTTAAACTGATGGCGGAACTC
GAGtaaTTTCCCGGGACCGTGTTGACAATTAATCATCGGCATAGTATATCGGCATAGTATAATACGACAATCAGATA
TACTGTCGTAAGGAGGTTGATTATGCCGGACGCTAAAAAGCAAGGCCGCTCTAATAAAGCTATGACCTTTTTTGT
ATGTTTTCTGGCTGCCCTTGCAGGGTTGCTTTTCGGCCTGGATATTGGGGTGATTGCCGGTGCTCTGCCCTTC
ATCGCTGATGAATTTCAAATTACTTCCCACACTCAAGAATGGGTAGTATCAAGTATGATGTTTGGCGCAGCGGTA
GGCGCAGTAGGTTCAGGCTGGTTAAGCTTCAAGTTGGGCCGCAAGAAGAGCCTGATGATCGGTGCTATTTTAT
TTGTCGCCGGGTCGTTGTTCTCGGCGGCAGCGCCGAACGTTGAAGTCCTTATCCTTAGTCGCGTGCTGCTGG
GTCTTGCGGTGGGCGTTGCATCATATACGGCGCCACTTTACCTGTCAGAAATTGCACCAGAAAAGATCCGTGG
AAGTATGATCTCCATGTATCAATTGATGATTACGATTGGAATTTTGGGCGCTTACTTGAGCGACACAGCTTTTTCA
TATACCGGAGCATGGCGTTGGATGTTAGGAGTTATCATTATCCCTGCGATTTTGTTGCTTATCGGAGTGTTCTTC
TTACCTGATTCTCCCCGTTGGTTTGCAGCGAAACGCCGTTTCGTCGACGCGGAACGTGTGCTGTTACGTTTAC
GTGACACGTCGGCAGAAGCTAAGCGTGAGTTAGATGAGATCCGCGAGTCATTGCAAGTCAAACAATCGGGGT
GGGCGCTTTTTAAGGAGAACAGCAACTTCCGTCGCGCCGTATTCCTGGGTGTGTTACTGCAGGTTATGCAACA
ATTCACTGGAATGAACGTGATTATGTATTATGCCCCAAAGATTTTTGAACTGGCTGGATACACCAATACTACCGAG
CAGATGTGGGGCACTGTCATTGTGGGTCTTACCAATGTCTTAGCTACTTTCATCGCAATTGGATTGGTAGATCGT
TGGGGTCGCAAGCCAACCCTGACTTTGGGATTTTTGGTTATGGCAGCGGGTATGGGTGTATTGGGTACGATGA
TGCATATCGGCATCCATAGTCCTAGCGCCCAATATTTTGCCATCGCTATGTTACTGATGTTCATTGTCGGCTTCG
CGATGAGTGCCGGACCGTTGATTTGGGTCTTATGCAGCGAGATCCAGCCATTGAAGGGCCGTGACTTTGGTAT
TACATGCTCAACAGCTACCAATTGGATTGCTAACATGATCGTTGGTGCGACCTTCTTGACGATGCTGAATACGTT
GGGCAACGCAAATACGTTTTGGGTGTACGCTGCTCTGAACGTCTTGTTCATCCTGCTTACTCTGTGGCTGGTC
CCGGAAACGAAGCACGTATCTTTGGAGCATATCGAGCGCAATTTGATGAAAGGGCGTAAGTTGCGTGAGATTG
GGGCTCATGATTAAAGATCGGCACGTAAGAGGTTCCAACTTTCACCATAATGAAATAAGATCACTACCGGGCGT
ATTTTTTGAGTTATCGAGATTTTCAGGAGCTAAGGAAGCTAAAATGAGGGAAGCGGTGATCGCCGAAGTATCGA
CTCAACTATCAGAGGTAGTTGGCGTCATCGAGCGCCATCTCGAACCGACGTTGCTGGCCGTACATTTGTACGG
CTCCGCAGTGGATGGCGGCCTGAAGCCACACAGTGATATTGATTTGCTGGTTACGGTGACCGTAAGGCTTGAT
GAAACAACGCGGCGAGCTTTGATCAACGACCTTTTGGAAACTTCGGCTTCCCCTGGAGAGAGCGAGATTCTC
CGCGCTGTAGAAGTCACCATTGTTGTGCACGACGACATCATTCCGTGGCGTTATCCAGCTAAGCGCGAACTGC
AATTTGGAGAATGGCAGCGCAATGACATTCTTGCAGGTATCTTCGAGCCAGCCACGATCGACATTGATCTGGCT
ATCTTGCTGACAAAAGCAAGAGAACATAGCGTTGCCTTGGTAGGTCCAGCGGCGGAGGAACTCTTTGATCCGG
TTCCTGAACAGGATCTATTTGAGGCGCTAAATGAAACCTTAACGCTATGGAACTCGCCGCCCGACTGGGCTGG
CGATGAGCGAAATGTAGTGCTTACGTTGTCCCGCATTTGGTACAGCGCAGTAACCGGCAAAATCGCGCCGAAG
GATGTCGCTGCCGACTGGGCAATGGAGCGCCTGCCGGCCCAGTATCAGCCCGTCATACTTGAAGCTAGACAG
GCTTATCTTGGACAAGAAGAAGATCGCTTGGCCTCGCGCGCAGATCAGTTGGAAGAATTTGTCCACTACGTGA
AAGGCGAGATCACCAAGGTAGTCGGCAAATAATTTTGCCGCAGTTTATGCTTCCGCCAGCGGCGGCAAAATCA
ATTCATCGCTCTCATGCTGCTGGGTGTAGCGCATCACTTCCAGTACGCGCAACCCCGCTCGGTGCACTGCATC
GGTTAACGCCTTCCCTTTCAGCAAGCCACTGATGAGCTGAGCACAAAA

**ΔgntR::EM7-gdh-gnl-zeoR**

GCGAGATGGTTCATTGAAAGTGCATCAGGATATGGAACTGTACCGCTGGGCGTTGCTGAAAGATGAGCAGTCG
GTGCATCAGATTGCCGCTGAACGCCGCGTCTGGATCCAGGTGGTGAAAGGCAATGTCACCATTAACGGCGTG
AAAGCCTCGACCAGCGATGGTCTGGCAATCTGGGATGAGCAGGCAATCTCCATCCATGCGGATAGCGACAGC
GAAGTGTTACTGTTCGATCTGCCGCCGGTTtaaCCGTGTTGACAATTAATCATCGGCATAGTATATCGGCATAGTA
TAATACGACAAGGTGAGGAACTAAACCATGTACTCCGACTTAGAGGGTAAGGTGGTCGTGATTACAGGATCGGC

| |
|---|
| TTCGGGTCTGGGCCGCGCAATGGGGGTTCGCTTTGCGGCAGAAAAAGCAAAGGTGGTTATTAACTATCGCTCG<br>CGTGAATCGGAGGCGAACGACGTTCTTGAGGAGATCAAGAAAGTTGGCGGAGAAGCGATTGCAGTGAAAGGA<br>GATGTTACGGTCGAATCGGACGTGGCTAATTTGATCCAGTCCGCAGTGAAGGAGTTTGGAACTTTAGATGTAAT<br>GATTAACAACGCGGGTATCGAGAACGCAGTCCCCAGCCATGAGATGCCTTTGGAGGACTGGAATCGTGTAATT<br>AACACGAATTTGACGGGCGCGTTTTTGGGCTCGCGTGAAGCGATCAAATACTTCGTAGAACACGACATCAAGG<br>GATCAGTAATTAACATGAGCTCAGTCCACGAAAAGATCCCCTGGCCTTTGTTCGTCCATTACGCAGCTAGTAAA<br>GGTGGAATGAAGTTAATGACCGAGACTCTGGCGTTAGAGTACGCGCCCAAGGGTATCCGTGTCAACAACATCG<br>GTCCCGGTGCGATCAACACACCAATTAACGCTGAGAAATTTGCGGATCCCAAGAAACGTGCAGATGTAGAGTC<br>AATGATCCCTATGGGATACATTGGAAAACCAGAGGAGATTGCCGCCGTGGCAACCTGGCTGGCATCCTCAGAA<br>GCGTCCTACGTGACCGGCATTACACTTTTCGCCGATGGAGGTATGACTCTTTATCCATCCTTTCAAGCGGGTCG<br>TTAAttgGGGAACGGACCTAAacggctagctcagtcctaggtacagtgctagctactagtgaaagaggagaaatactagATGGCCAAAAATA<br>ACATGAATGGTTCTACCATCGGTAAGATCACTAAATTCAGCCCACGTCTGGATGCGATCCTGGATGTTAGCACTC<br>CGATTGAAGTGATTGCTTCTGACATCCAGTGGTCTGAGGGCCCGGTATGGGTTAAAAACGGTAACTTCCTGCT<br>GTTCTCCGACCCGCCGGCGAACATTATGCGTAAATGGACGCCGGACGCGGGTGTCTCTATTTTCCTGAAGCCG<br>TCTGGTCATGCAGAGCCTATCCCGGCAGGTCAGTTTCGTGAACCGGGCAGCAACGGTATGAAGGTAGGTCCT<br>GATGGTAAGATTTGGGTAGCCGATTCCGGTACCCGCGCTATTATGAAGGTTGACCCGGTTACTCGCCAGCGTT<br>CTGTTGTGGTGGATAACTACAAAGGTAAGCGTTTCAACTCTCCGAACGACCTGTTCTTCTCTAAATCCGGCGCT<br>GTTTATTTCACTGACCCGCCGTATGGCCTGACCAACCTGGACGAATCTGATATTAAAGAAATGAACTATAACGGT<br>GTTTTCCGCCTGAGCCCGGACGGTCGTCTGGACCTGATCGAAGCGGGCCTGTCCCGTCCGAATGGCCTGGC<br>TCTGAGCCCAGATGAGACCAAACTGTATGTTAGCAATTCTGACCGCGCTAGCCCAAACATCTGGGTTTACAGCC<br>TGGATTCTAACGGTCTGCCGACTAGCCGTACTCTGCTGCGCAACTTTCGCAAAGAGTATTTCGACCAAGGTCT<br>GGCCGGTCTGCCGGACGGTATGAACATCGATAAACAGGGTAACCTGTTCGCTAGCGCACCGGGCGGTATCTAC<br>ATTTTTGCACCGGATGGCGAATGCCTGGGTCTGATCAGCGGTAACCCGGGCCAGCCGCTGTCCAACTGCTGT<br>TTTGGTGAAAAGGGTCAGACTCTGTTCATTTCCGCCTCTCATAACGTAGTACGCGTGCGTACCAAAACGTTTGG<br>CTAATTTGTTTATTTTTCTAAATACATTCAAATATGTATCCGCTCATGAGACAATAACCCTGATAAATGCTTCAATAA<br>TATTGAAAAAGGAAGAGTATGGCTAAACTGACGTCGGCCGTTCCAGTGCTTACTGCGCGTGATGTAGCGGGAG<br>CCGTAGAGTTTTGGACGGATCGTCTTGGGTTTAGTCGCGACTTTGTGGAAGATGACTTCGCAGGGGTTGTTCG<br>TGATGACGTCACACTGTTCATCAGTGCCGTACAGGATCAGGTTGTACCCGATAACACTCTTGCGTGGGTATGGG<br>TGCGTGGCCTGGATGAGTTATACGCCGAATGGTCCGAGGTAGTCAGCACAAACTTCCGCGACGCATCCGGGC<br>CCGCTATGACTGAGATCGGGGAACAACCGTGGGGACGTGAGTTTGCCTTACGTGACCCGGCGGGGAACTGC<br>GTCCACTTTGTGGCGGAGGAGCAGGACTAGAAACCaaagaggagaaatactagATGttgAGCACGACTAACCATGATCA<br>CCACATTTACGTCTTGATGGGCGTATCGGGCAGCGGCAAATCTGCGGTCGCCAGTGAAGTGGCGCATCAACTT<br>CATGCCGCGTTTCTTGATGGCGATTTCCTCCATCCACGGCGCAATATCGAAAAAATGGCGTCTGGCGAACCAC<br>TGAATGACGACGATCGCAAACCGTGGTTGCAGGCGCTGAACGACGCCGCGTTTGCTATGCAGCGCACTAATAA<br>AGTGTCGC |

**Table S4:** Oligos used for strain confirmations & sequencing

| **Oligo** | **Sequence** |
|---|---|
| udhA_conf_F | CAAAAGAGATTCTGGGTATTCACT |
| bsdR_intR | GAGCATGGTGATCTTCTCAGT |
| fabI_int_F | GCAAAATGCTGGCTCATTG |

| | |
|---|---|
| gentR_intR | GCGATGAATGTCTTACTACGGA |
| gltA_int_F | TATCATCCTGAAAGCGATGG |
| zeo_intR | ACTGAAGCCCAGACGATC |
| yhhW_intF | GGATGAGCAGGCAATCTCC |
| gdh_intF | GTGCGATCAACACACCAATT |
| gnl_intF | GTTTCAACTCTCCGAACGAC |
| gntK_intR | CCCATCAAGACGTAAATGTGG |
| ptsH_intF | GTCTGCACACCCGCCCTG |
| pdxK_intR | GAATTGATTTTGCCGCCGCTG |
| specR_intR | GAGTCGATACTTCGGCGATCAC |
| galP_intR | CTAACTCACGCTTAGCTTC |

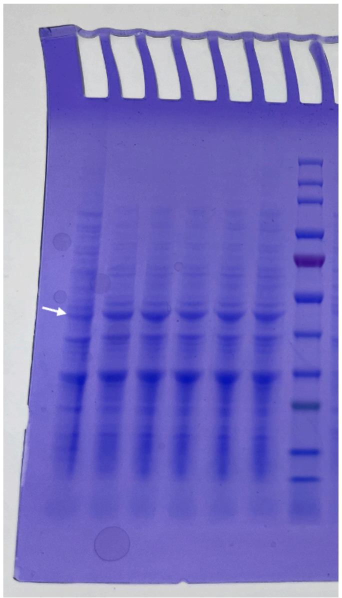

**Figure S1. Expression analysis of alanine dehydrogenase (Ald*) in different host strains.** Whole-cell lysates were analyzed by SDS–PAGE to assess Ald* expression. Densitometry indicates Ald* expression level of ~12% relative to total protein. Lane order (left to right):

DLFZ_25, DLF_GBP1, DLF_GBP3, DLF_GBP4, DLF_GBP7, DLF_GBP8, ladder. White arrow indicates the protein band corresponding to Ald*

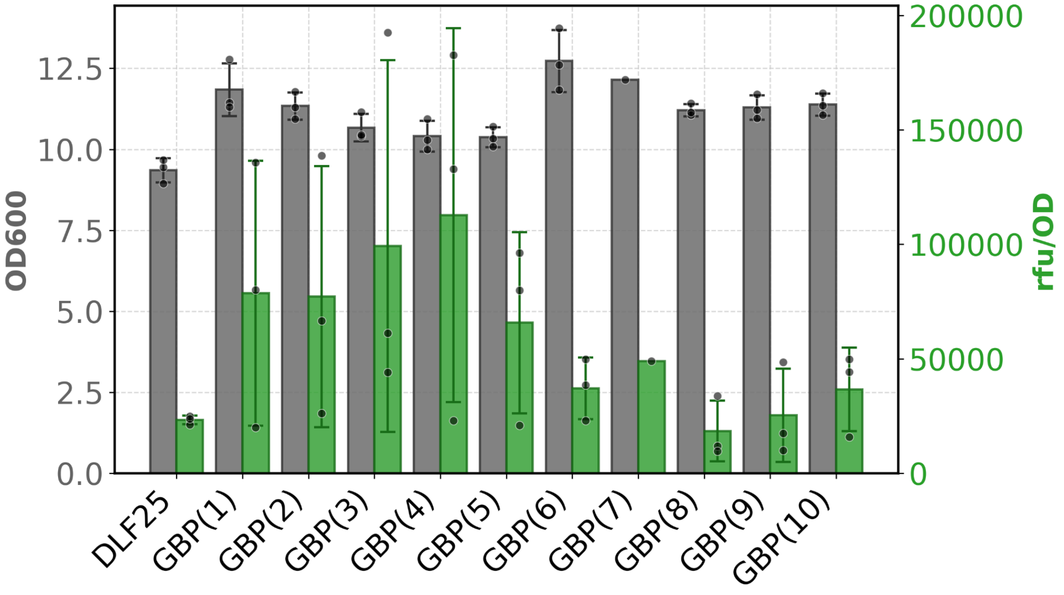


**Figure S2.** Growth profile and GFP expression of 10 colonies picked after construction of GBP strain

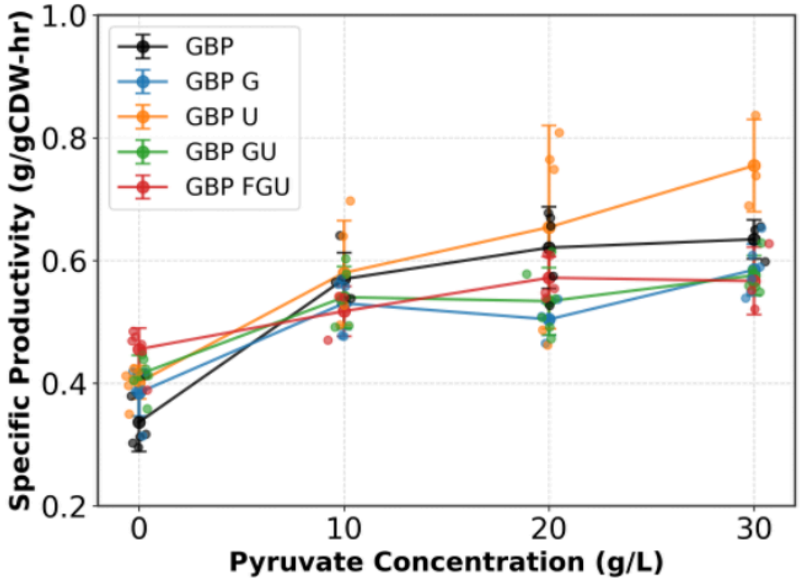


**Figure S3.** Alanine productivity of the best-performing strains with metabolic valves across different pyruvate concentrations

### References

Li, Shuai, Zhixia Ye, Juliana Lebeau, Eirik A. Moreb, and Michael D. Lynch. 2020. “Dynamic Control over Feedback Regulation Identifies Pyruvate-Ferredoxin Oxidoreductase as a Central Metabolic Enzyme in Stationary Phase E. Coli.” In *bioRxiv*. August 10. https://doi.org/10.1101/2020.07.26.219949.

Li, Shuai, Zhixia Ye, Eirik A. Moreb, et al. 2021. “Dynamic Control over Feedback Regulatory Mechanisms Improves NADPH Flux and Xylitol Biosynthesis in Engineered E. Coli.” *Metabolic Engineering* 64 (March): 26–40.

Menacho-Melgar, Romel, Zhixia Ye, Eirik A. Moreb, et al. 2020. “Scalable, Two-Stage, Autoinduction of Recombinant Protein Expression in E. Coli Utilizing Phosphate Depletion.” *Biotechnology and Bioengineering* 117 (9): 2715–2727.

Ye, Zhixia, Eirik A. Moreb, Shuai Li, et al. 2021. “Escherichia Coli Cas1/2 Endonuclease Complex Modifies Self-Targeting CRISPR/Cascade Spacers Reducing Silencing Guide Stability.” *ACS Synthetic Biology* 10 (1): 29–37.
